# Transcriptome Analysis Reveals Enhancement of Cardiogenesis-Related Signaling Pathways by S-nitroso-*N*-pivaloyl-D-penicillamine (SNPiP): Implications for Improved Diastolic Function and Cardiac Performance

**DOI:** 10.1101/2023.06.20.545771

**Authors:** Yasuhiro Takenaka, Masataka Hirasaki, Hidemasa Bono, Shigeo Nakamura, Yoshihiko Kakinuma

## Abstract

We previously reported a novel compound called S-nitroso-*N*-pivaloyl-D-penicillamine (SNPiP), which was screened from a group of nitric oxide (NO) donor compounds with a basic chemical structure of S-nitroso-*N*-acetylpenicillamine (SNAP), to activate the non-neuronal acetylcholine (NNA) system. SNPiP-treated mice exhibited improved cardiac output and enhanced diastolic function, without an increase in heart rate. The NNA-activating effects included increased resilience to ischemia, modulation of energy metabolism preference, and activation of angiogenesis. Here, we performed transcriptome analysis of SNPiP-treated mice ventricles to elucidate how SNPiP exerts beneficial effects on cardiac function. A time-course study (24 and 48 h after SNPiP administration) revealed that SNPiP initially induced Wnt and cGMP-protein kinase G (PKG) signaling pathways, along with upregulation of genes involved in cardiac muscle tissue development and oxytocin signaling pathway. We also observed enrichment of glycolysis-related genes in SNPiP-treated samples, indicating a metabolic shift from oxidative phosphorylation to glycolysis following SNPiP administration in the hearts. Additionally, SNPiP significantly upregulated atrial natriuretic peptide (ANP) and sarcolipin (SLN), which play crucial roles in calcium handling and cardiac performance. These findings suggest that SNPiP may have therapeutic potential based on the pleiotropic mechanisms elucidated in this study.

## Introduction

The non-neuronal acetylcholine (NNA) system, which is distinct from the parasympathetic nervous system (PNS), is a unique effector that influences local physiological functions in an autocrine/paracrine manner. The non-neuronal cardiac cholinergic system (NNCCS), which is a component of the NNA system, is present within cardiomyocytes and is responsible for the synthesis of acetylcholine (ACh) independently of the PNS [1–3]. Therefore, in the heart, ACh is derived from two sources: the PNS and the NNCCS. Using murine models with gain and loss of function, we and other groups have already reported that the NNCCS is a crucial system to sustain cardiac functions and provide resilience potential against hypoxia/ischemia by upregulating survival signals, modulating energy metabolism preference, enhancing angiogenesis, and activating cell–cell communication [4–7]. These findings prompted us to explore the development of an inducer or activator of the NNCCS. Based on the recognition that the NO donor SNAP exhibits relatively weak potency in inducing choline acetyltransferase (ChAT), we screened chemical compounds with a basic structure similar to that of SNAP.

NO is naturally produced in the body and has the ability to dilate blood vessels. NO also has many physiological functions, including regulation of blood pressure and protection of the heart, which has led to the development of NO-based therapies for heart disease [8, 9]. However, acute dilation of blood vessels can decrease blood pressure and concomitantly increase heart rate in a compensatory manner. Moreover, despite the effectiveness of NO-based drugs in the treatment of heart disease, their short half-lives make their use difficult. The half-lives of N-diazeniumdiolates (NONOates) is from 2 s to 20 h [10], whereas that of SNAP is approximately 6 h [11]. Therefore, the development of a novel NO-generating drug that causes sufficient and long-lasting NO release with less of an effect on hemodynamic parameters and that ultimately activates the NNCCS can provide a novel therapeutic modality that differs from conventional modalities.

We previously reported the effects of SNPiP, a novel NO donor and NNCCS activator, on cardiac function [12]. This compound releases NO for longer than traditional NO donors, indicating its potential to enhance cardiac function. Indeed, *in vivo* experiments with SNPiP-treated mice have demonstrated improved diastolic function without tachycardia, leading to enhanced cardiac output without an increase in heart rate which supports our hypothesis that the NNCCS plays a crucial role in sustaining cardiac functions, including potentiating resilience to ischemia, modulating energy metabolism preference, and activating angiogenesis [12]. SNPiP also improved impaired cardiac function in db/db hearts [13], suggesting its potential use as a therapeutic agent to activate the NNCCS and modulate cardiac function. It has been reported that SNPiP gradually elevates the intracellular levels of cyclic guanosine monophosphate (cGMP) and NO levels in H9c2 and HEK293 cells [12], which in turn upregulate ChAT and increase ACh synthesis. However, the detailed molecular mechanisms by which SNPiP affects cardiac function remain unclear.

To elucidate how SNPiP influences the expression of genes associated with cardiac function, we conducted a transcriptome analysis of ventricles from wild-type male mice treated with SNPiP. Gene Ontology (GO) analysis revealed that SNPiP upregulated genes involved in cardiac muscle development and cellular differentiation. Furthermore, the results of pathway analysis suggested that SNPiP activates the oxytocin signaling pathway, which is known to improve cardiac performance and stimulate cardiomyocyte differentiation. Lastly, we found that administration of SNPiP resulted in significant upregulation of atrial natriuretic peptide (ANP) and sarcolipin (SLN), which could be crucial for elucidating the comprehensive mechanisms of the effect of SNPiP on cardiac function. Our study provides a more in-depth understanding of the molecular mechanisms underlying the therapeutic effects of NO-based drugs and their potential adverse effects.

## Materials and Methods

### Mice and treatments

SNPiP, which has a molecular weight of 262.32, was synthesized using a previously reported method [12]. This compound, identified by product code: 197-19151, is now commercially available and can be procured from FUJIFILM Wako Chemicals (Osaka, Japan). Adult male C57/BL6 mice aged 12 weeks were used in the experiments. SNPiP was dissolved in dimethyl sulfoxide (DMSO) (D2650, Sigma-Aldrich Japan, Tokyo, Japan) and administered by intraperitoneal injection at a dose of 60 pmol/g body weight. As a control, a comparable dilution of the solvent was administered in the same manner [12]. After an appropriate amount of time, the heart was removed from a mouse euthanized by cervical dislocation. The atria were subsequently dissected from the heart, leaving the ventricles, which were then used as samples. RNA sequencing (RNA-seq) analysis

Total RNA was isolated from the whole ventricle of mice using ISOGEN reagent (Nippon Gene, Tokyo, Japan) according to the manufacturer’s instructions. The yield and quality of the purified RNA were evaluated by spectrophotometry, agarose gel electrophoresis, and Bioanalyzer assessments. The library preparation and RNA-seq were conducted by Macrogen Japan (Tokyo, Japan) using a TruSeq^TM^ Stranded Total RNA Library Prep Kit with a Ribo-Zero^TM^ kit and NovaSeq6000 paired-end sequencing (Illumina). The sequencing reads were aggregated into an rRNA reference (Mouse_rRNA_Reference_BK000964) using Bowtie package to remove rRNA reads. Unpaired reads were removed using FASTQ-pair. Paired reads were aligned using STAR version 2.7.8a with the GRCm38 reference alignment. RSEM was used to quantify the transcript. Transcripts Per Million (TPM) calculated using RSEM were log_2_(count+1) transformed.

Significance estimates of differences in gene expression, such as the *p*-value and False Discovery Rate (FDR), were calculated from expected counts using edgeR package in R. Differentially expressed genes (DEGs) were defined as genes with a *p*-value < 0.05 and |log_2_(fold change)| > 0.5. Using the count data, the Trimmed Means of M values (TMM) were calculated using the edgeR package. The principal component analysis (PCA), heatmap, dotplot, and volcano plot figures were generated using R version 4.2.3.

### Gene Ontology and pathway analysis

To characterize the molecular and functional aspects of the DEGs, GO and Kyoto Encyclopedia of Genes and Genomes (KEGG) pathway analyses were performed using the online DAVID database (https://david.ncifcrf.gov/). Pathway and network analyses were performed using QIAGEN Ingenuity Pathway Analysis (IPA). The DEGs were subjected to core analysis, which led to enrichment analysis of canonical pathways, prediction of diseases and biological functions, and identification of upstream regulators. The results of the core analysis for the 24 and 48 h samples were visualized as heatmaps using the Comparison Analysis feature of IPA. Gene set enrichment analysis (GSEA)

Using TMM values, which are normalized count data, GSEA was performed according to the methods described on the GSEA website (http://www.gsea-msigdb.org/gsea/index.jsp).

### Western blot analysis

Tissue lysates were prepared from cardiac ventricles excised from the heart using T-PER^TM^ Tissue Protein Extraction Reagent (ThermoFisher Scientific, Tokyo, Japan) according to the manufacturer’s instructions. After centrifugation, the tissue lysate supernatant was collected and treated with SDS sample buffer. Comparable amounts of protein (10–30 μg/lane), quantified using a BCA kit, were loaded (as subsequently verified based on the level of GAPDH). Tissue lysate supernatants were electrophoresed and blotted onto polyvinylidene fluoride membranes. After blocking with 4% skim milk, the membranes were incubated with the following primary antibodies: a rabbit polyclonal antibody against ANP (N-20, 1:500, Santa Cruz Biotechnology, Dallas, TX, USA), sarcolipin (ABT13, 1:1000, MERCK KGAA, Darmstadt, Germany), or a rabbit monoclonal antibody against GAPDH (1:5000, Cell Signaling Technology, Danvers, MA, USA), followed by reaction with a secondary antibody conjugated with horseradish peroxidase (HRP). The antigen-HRP signals were detected using ImmunoStar LD reagent (FUJIFILM Wako Chemicals) and a C-DiGit Blot Scanner (LI-COR Corp., Lincoln, NE, USA).

## Results

### Transcriptome analysis of SNPiP-administered ventricles of wild-type mice

Male wild-type C57/BL6 mice were intraperitoneally administered with SNPiP dissolved in DMSO (Fig. 1A). Male mice were used to exclude sexual dimorphism particularly the beneficial effects of estrogen on cardiovascular diseases. The time points for sacrificing SNPiP-treated mice (24 and 48 h) were determined based on our previous studies. In these studies, the effect of SNPiP on cardiac dilation was most pronounced 48 and 72 h after administration [12, 13]. To investigate how SNPiP affects the expression of genes related to cardiac function, changes in gene expression in wild-type mice treated with DMSO (control) or SNPiP were determined by RNA-seq.

**Figure 1.**
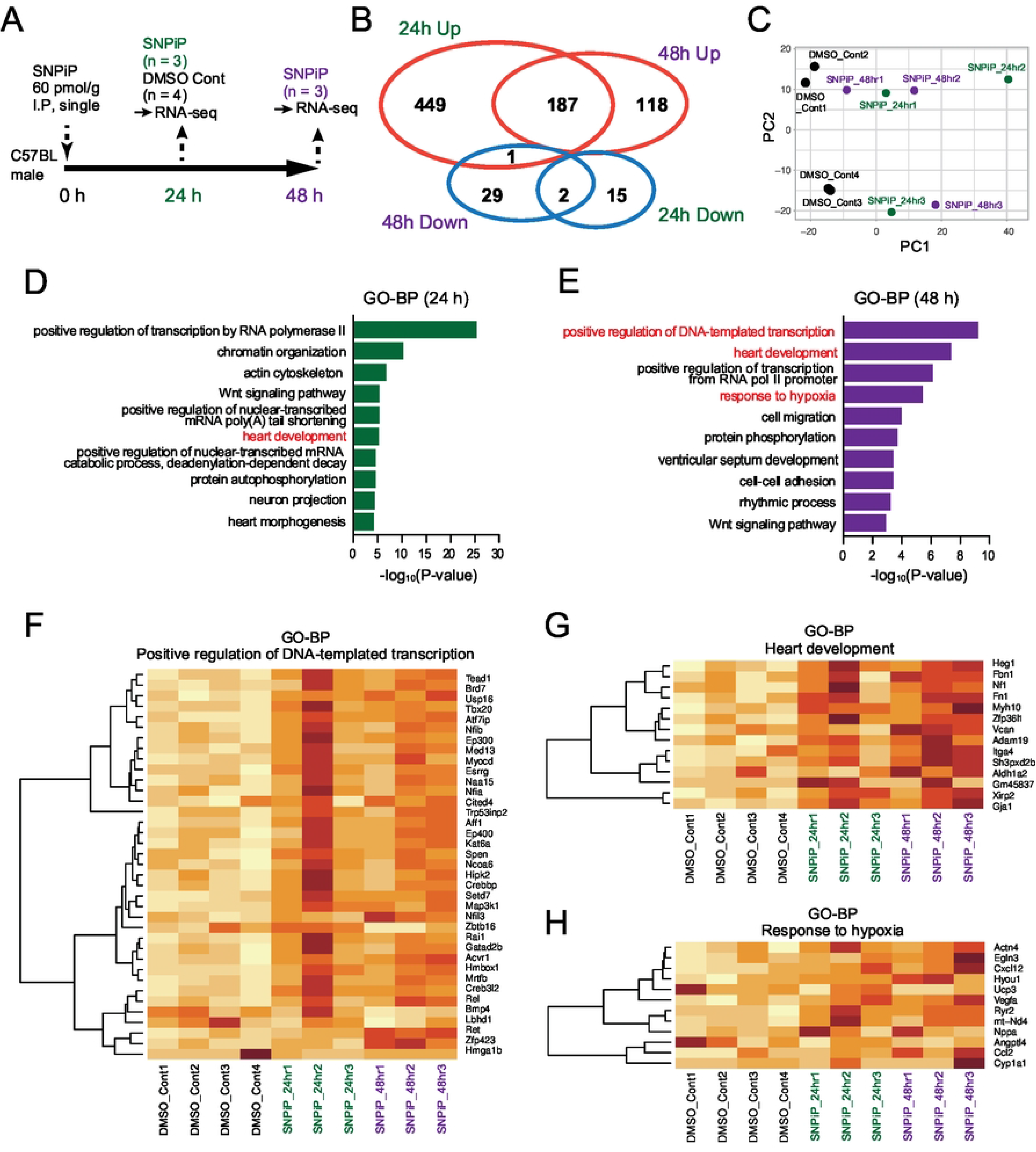
(A) The experimental design for SNPiP treatment of wild-type C57BL male mice. Whole ventricles were isolated at 24 and 48 h after intraperitoneal (I.P) administration of SNPiP. (B) Venn diagram displaying the number of upregulated and downregulated genes at 24 and 48 h after SNPiP administration, respectively (*p* < 0.05, |log_2_(fold change)| > 1). (C) Principal component analysis of TMM-normalized reads in DMSO controls (n = 4), SNPiP at 24 h (n = 3), and SNPiP at 48 h (n = 3). The contribution rate of the first two principal components (PC1 and PC2) is 53.6%. Summary of biological process in gene ontology analysis (GO-BP) for upregulated genes via SNPiP administration at 24 h (D) and 48 h (E) samples. Heatmap of genes differentially expressed in the GO-BP terms “positive regulation of DNA-templated transcription” (F), “heart development” (G), and “response to hypoxia” (H).

We found that SNPiP upregulated 636 genes (24 h) and 305 genes (48 h) with a log_2_ fold change (FC) greater than 0.5 (*p* < 0.05), whereas, it downregulated 17 genes (24 h) and 32 genes (48 h) with a log_2_FC less than −0.5 (*p* < 0.05), indicating predominant upregulation of genes following SNPiP treatment (Fig. 1B). A total of 187 genes were upregulated between 24 and 48 h, whereas two genes were downregulated. Additionally, only one gene (Tob2, ENSMUSG00000048546) exhibited counterregulation between 24 and 48 h after SNPiP administration.

We conducted PCA to visualize the differences and similarities between the samples (Fig. 1C). Plotting of the data onto the first two principal components revealed that the SNPiP-treated samples exhibited better segregation from the DMSO controls in the first principal component (PC1). However, biological replicates were divided into two groups in the second principal component (PC2), indicating substantial variation in gene expression following SNPiP administration among individuals.

GO analysis using the DAVID database revealed that many of the upregulated genes were enriched for positive regulation of transcription and heart development-related processes at 24 h (Fig. 1D) and 48 h (Fig. 1E), suggesting that SNPiP caused regulation of transcription or heart development. Heatmap analysis revealed pronounced upregulation of genes in the GO biological process (BP) terms “positive regulation of DNA-templated transcription” (Fig. 1F) and “heart development” (Fig. 1G) in three biological replicates at both 24 and 48 h. We also identified enrichment of the GO-BP term “response to hypoxia” at 48 h (Fig. 1E) and differential expression of crucial genes in hypoxia signaling such as *Egln3*, *Vegfa*, *Nppa*, and *Angptl4* (Fig. 1H).

### Gene Set Enrichment Analysis (GSEA)

Next, we employed GSEA and found enrichment of the mitochondrial ATP biosynthetic process in the DMSO controls (Figs. 2A and 2B), which indicated a reduction in respiratory activity in SNPiP-treated samples. Indeed, many cytochrome c oxidase genes in complex IV of the mitochondrial electron transport chain were downregulated in the SNPiP-treated ventricles (Fig. 2C). Whereas, we identified enrichment of the glycolytic process in SNPiP-treated samples (Figs. 2D-2F), suggesting a metabolic shift from oxidative phosphorylation to the glycolytic process by administration of SNPiP in mouse hearts. In particular, genes induced by SNPiP were involved in insulin (*Insr* and *Igf1*), mTOR (*mTor* and *Rptor*), and AMPK (*Prkaa2*) signaling pathways (Fig. 2F). Furthermore, the regulation of smooth muscle cell differentiation appeared to be suppressed in the SNPiP-treated samples (Figs. 2D and 2E), implying that the differentiation of coronary artery smooth muscle cells was inhibited by SNPiP treatment.

**Figure 2.**
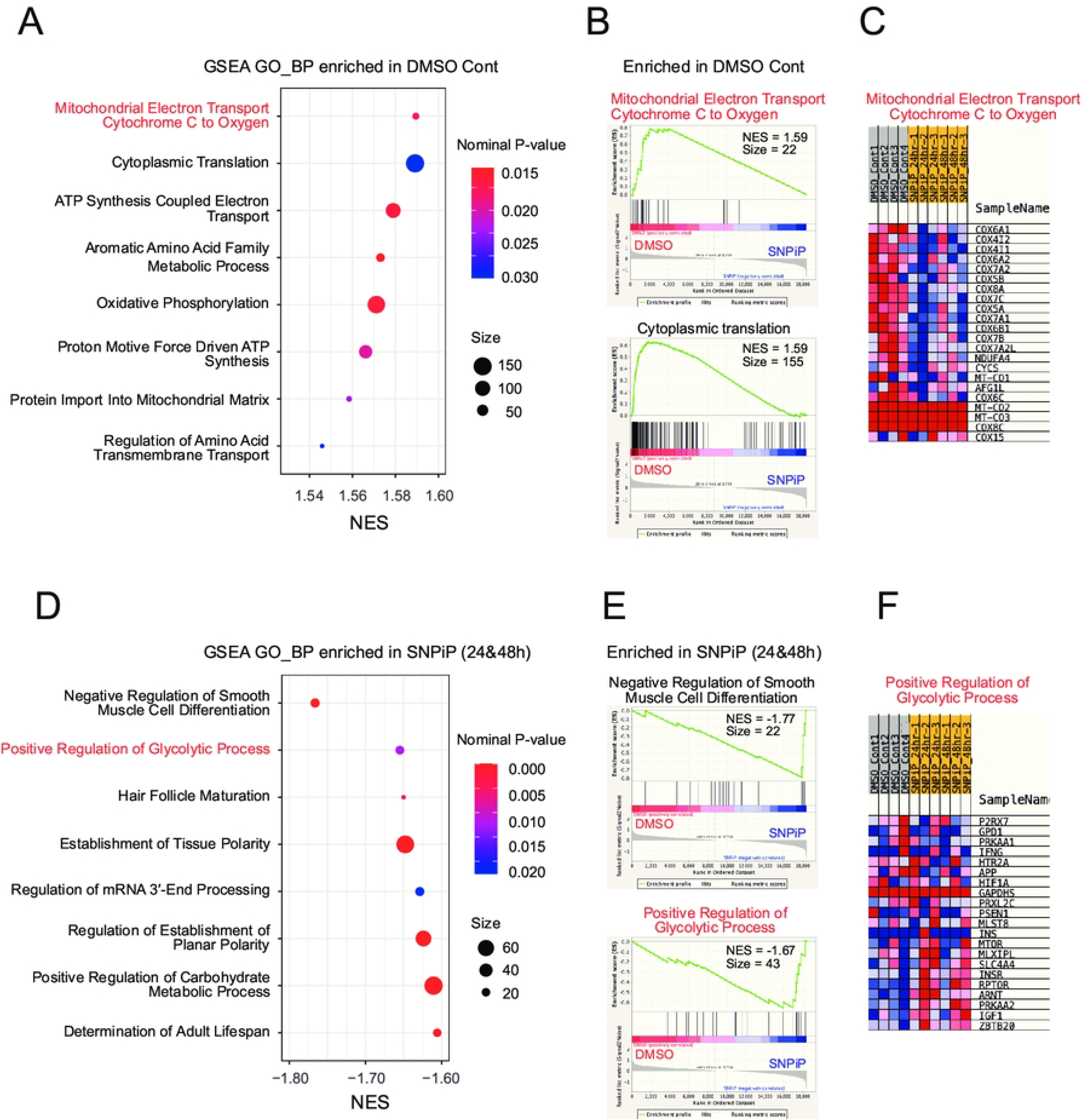
(A) Summary of the GO-BP terms enriched in DMSO controls by GSEA analysis. NES, normalized enrichment score. (B) GSEA plots of the top two-ranked GO-BP enriched in DMSO controls. (C) Heatmap of differentially expressed genes in the GO-BP term “mitochondrial electron transport cytochrome c to oxygen”. (D) Summary of GO-BP terms enriched in SNPiP-treated samples at 24 and 48 h by GSEA analysis. (E) GSEA plots of the top two-ranked GO-BP enriched in SNPiP-treated samples. (F) Heatmap of differentially expressed genes in the GO-BP term “positive regulation of glycolytic process”.

### Pathway analyses

KEGG pathway analysis revealed that enriched pathways mainly involved “Wnt signaling pathway” and “cGMP-PKG signaling pathway” were remarkably affected by the administration of SNPiP at 24 h (Fig. 3A), compared with the DMSO controls. Figure 3B illustrates KEGG pathway mapping for “Wnt signaling pathway”. More than 10 genes in the canonical Wnt pathway were upregulated, while six genes in the Wnt/Ca^2+^ pathway were also identified (Fig. 3B), implying that SNPiP triggered Ca^2+^ signaling, which may be involved in regulating cardiac contraction and dilation. The analysis also identified that “oxytocin signaling” was enriched at both 24 and 48 h (Fig. 3A). Ingenuity pathway analysis (IPA) also revealed significant activation of multiple signaling pathways, including the “Wnt/Ca^2+^ pathway” and “oxytocin signaling pathway” in addition to other molecular signaling pathways such as “estrogen receptor signaling”, particularly at 24 h after SNPiP administration (Fig. 3C). Differential gene expression in the “oxytocin signaling pathway” was visualized by hierarchical clustering heatmap analysis (Fig. 3D), and the IPA pathway map (Fig. 3E). Although the induction levels varied among biological replicates, the overall gene expression in the oxytocin signaling pathway was elevated upon SNPiP treatment. Analysis of IPA diseases and bio-functions revealed that SNPiP treatment inhibited hypoplasia and apoptosis of the heart, and activated cardiogenesis at both 24 and 48 h (Fig. 3F), thus supporting the results of the GO-BP analyses (Figs. 1D, 1E, and 1G). Upstream regulator analysis using IPA predicted several potential upstream regulators (Fig. 3G). Some of these candidate genes exhibited differential expression, i.e., *Myocd*, with an average log_2_FC of 0.82 at 24 h, *Mrtfa*, with a log_2_FC of 0.75 at 24 h, and *Mrtfb*, with a log_2_FC of 0.65 at 24 h post SNPiP administration. However, further investigation is required to elucidate the protein expression levels and detailed relationships between these upstream regulators and downstream effectors.

**Figure 3.**
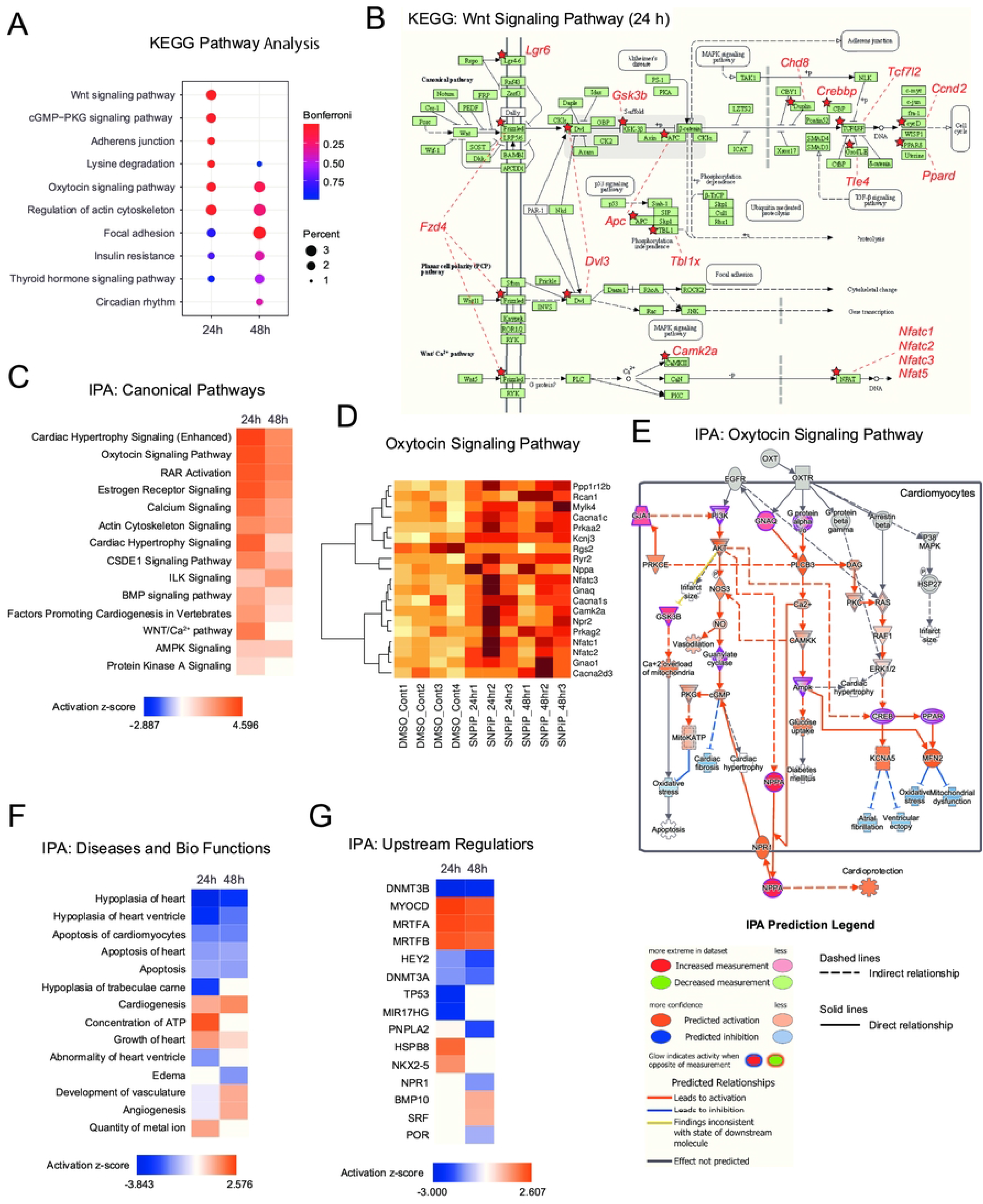
(A) Summary of the KEGG pathway analysis of differentially expressed genes between SNPiP-treated and control (DMSO-treated) samples. (B) Mapping of the differentially expressed genes in the KEGG Wnt signaling pathway. Genes exhibiting upregulation (*p* < 0.05, |log_2_(fold change)| > 1) are denoted in red font. (C) Summary of the IPA canonical pathway analysis of differentially expressed genes between SNPiP-treated and control samples. The activation z-score provides an indication of the probable activation states of a pathway. (D) Heatmap of differentially expressed genes related to the “oxytocin signaling pathway” in the KEGG pathway. (E) IPA pathway map of oxytocin signaling in cardiomyocytes. The legends for IPA predictions displayed on the map are provided below. (F) Heatmap representing diseases and biological functions predicted by IPA. Activation z-score: red indicates activation, blue indicates inactivation. (G) Heatmap representing upstream regulators predicted by IPA.

### Volcano plots and network analysis

We identified highly upregulated (log_2_FC > 1) and downregulated (log_2_FC < −1) genes by SNPiP at 24 and 48 h (*p* < 0.05), some of which were common to both time points (Figs. 4A and 4B). These shared genes play important roles in the response to hypoxia, tissue development, Ca^2+^ homeostasis, and the regulation of muscle relaxation (Table 1). Among them, *Nppa* encodes the cardiac hormone ANP, which promotes vasodilation, reduces heart workload and lowers blood pressure. We confirmed the significant upregulation of ANP levels in the SNPiP-treated murine ventricles by immunoblotting (Fig. 4C). Similarly, *Sln* encodes SLN, which plays a significant role in cardiac function, specifically in the regulation of Ca^2+^ handling by interaction with the sarcoplasmic/endoplasmic reticulum ATPase (SERCA) pump in cardiomyocytes. Upregulation of SLN levels by SNPiP was also evaluated by immunoblotting (Fig. 4D). Lastly, we also identified the protein–protein interaction networks associated with highly differentially expressed transcripts using STRING analysis. While many interactions were attributed to the co-expression of genes in publicly available gene expression datasets, physical protein–protein interactions (indicated by purple lines in Fig. 4E) were observed within the network between MYL7 and MYL4, suggesting enhanced cardiac muscle growth or remodeling processes by SNPiP.

**Figure 4.**
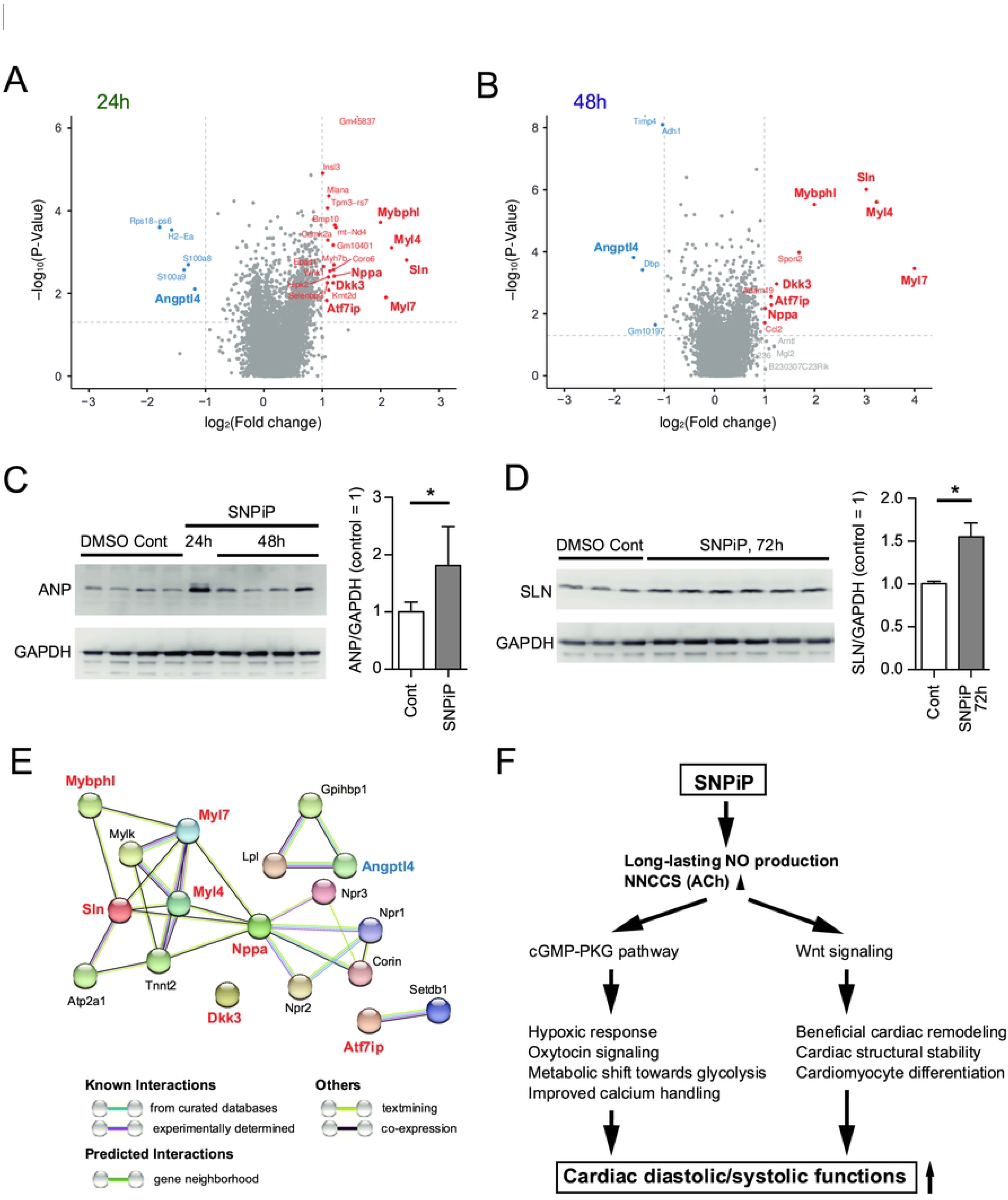
Volcano plots of differentially expressed genes between controls (DMSO-treated) and SNPiP-treated samples at 24 h (A) and 48 h (B). Gene names are displayed for highly upregulated (red) and downregulated (blue) genes (*p* < 0.05, |log_2_(fold change)| > 1). Genes that are consistently upregulated or downregulated at both 24 and 48 h following SNPiP administration are highlighted in bold font. Immunoblot analysis of ANP (*Nppa*) (C) and SLN (*Sln*) (D) in samples treated with DMSO or SNPiP. The asterisk (*) indicates *p* < 0.05 by one-tailed unpaired *t*-test. (E) STRING network analysis of highly upregulated (red) and downregulated (blue) genes. (F) Schematic representation of the pathways and biological functions through which SNPiP enhances cardiac diastolic function.

**Table 1.** Highly upregulated and downregulated genes by SNPiP at 24 and 48 h

| **Ensembl gene ID** | **Gene symbol** | **Gene name** | **log<sub>2</sub>FC** | **p-Value** | **FDR** |
| --- | --- | --- | --- | --- | --- |
| ENSMUSG00000020469 | *Myl7* | myosin, light polypeptide 7, regulatory | 3.046 | 5.189.E-03 | 0.287 |
| ENSMUSG00000042045 | *Sln* | sarcolipin | 2.739 | 5.973.E-04 | 0.161 |
| ENSMUSG00000061086 | *Myl4* | myosin, light polypeptide 4 | 2.715 | 3.326.E-04 | 0.144 |
| ENSMUSG00000068745 | *Mybphl* | myosin binding protein H-like | 1.996 | 1.757.E-04 | 0.102 |
| ENSMUSG00000030772 | *Dkk3* | dickkopf WNT signaling pathway inhibitor 3 | 1.216 | 4.418.E-03 | 0.271 |
| ENSMUSG00000041616 | *Nppa* | natriuretic peptide type A | 1.108 | 1.063.E-02 | 0.341 |
| ENSMUSG00000030213 | *Atf7ip* | activating transcription factor 7 interacting protein | 1.107 | 2.153.E-03 | 0.208 |
| ENSMUSG00000002289 | *Angptl4* | angiopoietin-like 4 | -1.401 | 3.828.E-05 | 0.067 |

## Discussion

In this report, we show the time-course effect of a novel NO donor, SNPiP, on the cardiac gene expression profile of wild-type mouse ventricles. In SNPiP-treated mice, diastolic function was greatly enhanced, suggesting increased left ventricular compliance, resulting in improved cardiac output, probably through the Frank–Starling law; hence, the drug potentially provides a new treatment strategy for cardiovascular diseases. Although the specific effects of SNPiP on protein levels, modification, and interaction remain to be elucidated, our transcriptome analysis revealed that SNPiP treatment promotes heart tissue development and cardiomyocyte differentiation. The terms annotated in the enrichment analysis primarily relate to embryonic heart development. However, the adult heart is capable of repair and regeneration [14]. Therefore, they can also be applied to certain aspects of heart repair and regeneration, along with the maintenance of proper cardiac function in adult organisms by maintaining the structural and mechanical integrity of the cardiomyocytes. An exogenous NO donor, DETA-NONOate, promotes the differentiation of Sca-1^+^ cardiac progenitor cells (CPC) towards the cardiac myocyte lineage with upregulation of cardiac structural genes such as *TnnT2* and *Myh7* [15]. Another NO donor, SNAP, also promotes cardiac differentiation in mouse embryonic stem (ES) cells [16]. Thus, we hypothesized that the NO produced by SNPiP triggers cardiogenesis and beneficial remodeling in the heart, resulting in improved cardiac function (Fig. 4F). Moreover, we previously reported that SNPiP induces NNCCS, leading to a gradual increase in ACh synthesis in cardiomyocytes [8]. This in turn upregulates NO production in an autocrine/paracrine manner, and NO from the donor alternatively upregulates the hypoxia-responsive pathway, as described later [17, 18]. Taken together, the enhanced NO production from SNPiP and SNPiP-induced NNCCS can synergistically promote mechanical stability through upregulation of developmentally critical machinery.

In our KEGG pathway analysis, genes in the Wnt signaling pathway were notably enriched following SNPiP administration (Figs. 3A and 3B). Wnt signaling has biphasic and stage-specific effects on cardiogenesis and hematopoiesis during mouse ES cell differentiation. Specifically, Wnt activation promotes differentiation into the cardiac lineage during the early phase but inhibits cardiomyogenesis during the late phase [19]. DETA-NONOate inhibits the canonical Wnt/β-catenin signaling pathway in Sca-1^+^ CPCs, which is required for induction of cardiac differentiation towards the cardiac myocyte lineage [15]. Given the enrichment of genes associated with tissue development by GO-BP analysis, activation of the Wnt signaling pathway via SNPiP administration may be relevant to cardiogenesis.

KEGG and IPA pathway analyses further demonstrated that SNPiP induced upregulation of genes within the oxytocin signaling pathway (Figs. 3A, 3C-3E). The oxytocin receptor (OXTR) is not only expressed in the uterus, mammary glands, and the brain but also in the heart of rodents [20, 21]. Oxytocin plays a crucial role in regulating cardiac function in addition to facilitating parturition and lactation. Specifically, oxytocin provides cardioprotection through various mechanisms, including negative chronotropic and inotropic effects by releasing ANP [22], exhibiting anti-inflammatory and antioxidant properties [23], and regulating glucose utilization [24]. Oxytocin also promotes the differentiation of cardiomyocytes, endothelial cells, and smooth muscle cells through NO release, which enhances cardiac development and function [25]. As SNPiP activated the expression of many genes involved in cardiac development (Fig. 1G) and glycolytic process (Figs. 2D-2F), it is tempting to speculate that this upregulation may be directly or indirectly mediated by the oxytocin signaling pathway. This hypothesis of oxytocin involvement is supported by the volcano plot analysis of transcriptome data and immunoblot analysis, which revealed remarkable upregulation of *Nppa*, the gene encoding ANP, one of the target genes of oxytocin signaling, at both 24 and 48 h post-drug administration (Figs. 4A and 4B), as well as a significant increase in ANP levels (Fig. 4C). Further investigations are needed to clarify the relationship between SNPiP and oxytocin signaling in cardiac function.

SLN, a small regulatory protein that negatively modulates the function of the SERCA pump to maintain intracellular Ca^2+^ levels within a physiological range, was also significantly upregulated at both the transcript and protein levels following SNPiP treatment (Figs. 4A, 4B, and 4D). In the *Sln* knockout (KO) model, enhanced SERCA activity results in abnormal intracellular Ca^2+^ handling and atrial remodeling, suggesting that dynamic regulation of the SERCA pump by SLN is crucial for maintaining cardiac contractility under normal and pathophysiological conditions [26, 27]. SLN also affects muscle function by signaling for muscle remodeling because Ca^2+^ acts as a potent cell signaling molecule. In skeletal muscle, high SLN levels increase oxidative metabolism and exercise capacity [28]. In contrast, upregulation of SLN in several different cardiac pathologies has been reported [29, 30]. In the left ventricles of patients with mitral regurgitation, the SLN protein level has been reported to be six-fold higher than that in normal controls [29]. Furthermore, adenoviral overexpression of SLN in adult rat ventricles resulted in decreased myocyte contractility and calcium transients [31]. Upregulation of SLN in pathological cardiac conditions, which are in an energy-starved state, may be due to compensation for the increased energy demand and sustained cardiac contraction. Therefore, a properly balanced level of SLN in cardiomyocytes should be indispensable for cardiac homeostasis. Thus, the modest upregulation of SLN protein (~1.6-fold) by SNPiP should be interpreted as being within the physiological range or an indication of pathological effects.

In addition to *Nppa* and *Sln*, we identified several genes that are highly upregulated following SNPiP administration (Figs. 4A and 4B). Intriguingly, four of these genes (*Nppa*, *Sln, Myl4*, and *Mybphl*) are upregulated in the left ventricles of endothelin receptor type B (ET_B_) KO (*EdnrB*^−/+^) mice but downregulated in wild-type (*EdnrB*^+/+^) mice under hypoxic and normoxic conditions [32]. Although *EdnrB* expression levels were not altered by SNPiP administration (data not shown), we believe that the similarities in the gene expression profiles between our model and the ET_B_ receptor study imply a common mechanism influencing cardiac function. Indeed, GO analysis revealed enrichment in the BP term “response to hypoxia” in SNPiP-treated ventricles at 48 h (*p* = 3.83 × 10^-6^) (Fig. 1E). The significant differential regulation of four key genes involved in hypoxia signaling *Egln3*, *Vegfa*, *Nppa*, and *Angptl4* (Fig. 1H), which are among the top 25 genes with high hypoxia-normoxia scores [33], supports the hypothesis that SNPiP induces hypoxic conditions. One of the upregulated genes, *Egln3* (Phd3), encodes prolyl 4-hydroxylase domain protein (PHD) and plays a crucial role in the hypoxia-inducible factor (HIF)-regulatory feedback loop [34]. Phd3 is directly induced by HIF [35] and hypoxic conditions [36] to compensate for the reduced oxygen availability. Noteworthy in this regard is the fact that NO can inhibit PHD activity, nevertheless in turn, inhibition of PHD by NO leads to HIF-1α accumulation, even under normoxic conditions [18, 37]. ANP is also regulated by HIF-1α and interacts with PKG, thereby converting it into a common pathway involving NO and the NNCCS. Thus, SNPiP-derived NO may inhibit PHD activity and cause a hypoxic response, although changes in HIF levels by SNPiP remain to be investigated. These findings suggest a potential molecular link between our model and the ET_B_ KO model within the downstream pathway of the hypoxia response.

Of note, the top-scored network among counterregulated genes, including *Nppa*, *Sln*, and *Myl4*, in *EdnrB*^+/+^ and *EdnrB*^−/+^ mice is related to the cGMP metabolic process and its connection to Ca^2+^ homeostasis [32]. NO upregulates cGMP, the second messenger of NO, in the heart, which activates PKG and eventually regulates Ca^2+^ homeostasis by either enhancing Ca^2+^ release via ryanodine receptors from the sarcoplasmic reticulum or by inhibiting L-type calcium channel [38]. We found that SNPiP increased cGMP levels in HEK293 cells [12] and activated the cGMP-PKG signaling pathway *in vivo* (Fig. 3A). Therefore, the regulation of Ca^2+^ homeostasis through cGMP-PKG signaling appears to be another mechanism shared between our model and the ET_B_ receptor study. However, the ET_B_ receptor plays a crucial role in mediating NO production [39], and in a blood flow cessation model of *EdnrB* KO mice, the level of NOx, a metabolite of NO, in the carotid artery was significantly lower compared to that of WT mice [40]. Therefore, ET_B_ receptor-mediated NO release may be reduced and ANP can stimulate the production of cGMP in an autocrine/paracrine manner [41] in *EdnrB*^−/+^ mouse ventricles. Since only limited information regarding NO and NOS levels in the hearts of *EdnrB^−/+^* mice is available, further investigation is warranted to address this issue and provide a more comprehensive level of understanding. Finally, with regard to *Myl4*, it is worth noting that previous studies have reported that loss-of-function models of *Myl4* exhibit morphological and electrical remodeling, leading to the development of arrhythmias [42, 43]. Therefore, SNPiP may play a protective role in cardiac functions by upregulating *Myl4* to suppress these remodeling processes.

Our previous studies demonstrated that SNPiP possesses the characteristic advantage of inotropic action by enhancing cardiac dilatation without inducing tachycardia [12]. This is in contrast to conventional inotropic agents, which enhance oxygen consumption and require more energy substrate resulting in a substantial cardiac workload. Furthermore, the mode of action of SNPiP is expected to be indolent based on the production mode of cGMP and NO consequently enhancing ACh production through NNA and NNCCS. This is supported by the observation that SNPiP enhances murine cardiac function to its maximum extent within 48 to 72 h [12]. As shown in this study, SNPiP triggers the expression of various genes involved in multilateral functions, and *de novo* protein synthesis induced by SNPiP may be integrated to perform such a cardioprotective action. In contrast, it is important to note that SNPiP is administered intravenously rather than orally. Consequently, exploring the potential of modifying SNPiP for oral delivery could offer significant advantages for future clinical applications.

## Acknowledgements

We are grateful to Ms. Yuko Kai and Ms. Shino Oikawa for their valuable technical supports. We would like to express my sincere gratitude to ChatGPT (GPT-4), developed by OpenAI, for its invaluable assistance in editing the English text. We would like to thank Editage (www.editage.jp) for English language editing. The map of the Wnt signaling pathway in *Mus musculus* (mmu04310) was obtained from the KEGG (Kyoto Encyclopedia of Genes and Genomes) database, and we gratefully acknowledge their valuable resource.

## Funding

This work was partly supported by the Japan Society for the Promotion of Science Grants-in-Aid for Scientific Research (JSPS KAKENHI), Grant Number 16K08560, the Smoking Research Foundation, and The Vehicle Racing Commemorative Foundation.

## Author contributions

Designed the experiments: YK, Performed the experiments: YT, SN, and YK, Analyzed the data: MH and HB, Wrote the paper: YT, MH, HB, and YK

## Notes

### Competing Interest Statement

The authors have declared no competing interest.

